# npInv: accurate detection and genotyping of inversions mediated by non-allelic homologous recombination using long read sub-alignment

**DOI:** 10.1101/178103

**Authors:** Haojing Shao, Devika Ganesamoorthy, Tania Duarte, Minh Duc Cao, Clive Hoggart, Lachlan J.M. Coin

**Affiliations:** Institute for Molecular Bioscience, University of Queensland, St Lucia, Brisbane, QLD 4072 Australia; Department of Medicine, Imperial College London, London, SW7 2AZ, United Kingdom

**Keywords:** Inversion, non allelic homologous recombination(NAHR), inverted repeat(IR), split read, long read sequencing, NA12878

## Abstract

Detection of genomic inversions remains challenging. Many existing methods primarily target inversions with a non repetitive breakpoint, leaving inverted repeat (IR) mediated non-allelic homologous recombination (NAHR) inversions largely unexplored. We present npInv, a novel tool specifically for detecting and genotyping NAHR inversion using long read sub-alignment of long read sequencing data. We use npInv to generate a whole-genome inversion map for NA12878 consisting of 30 NAHR inversions (of which 15 are novel), including all previously known NAHR mediated inversions in NA12878 with flanking IR less than 7kb. Our genotyping accuracy on this dataset was 94%. We used PCR to confirm presence of two of these novel NAHR inversions. We show that there is a near linear relationship between the length of flanking IR and the size of the NAHR inversion.

## Background

Inversion polymorphisms, in which the orientation of a segment of DNA is flipped with respect to its ancestral orientation relative to the rest of the chromosome, were originally discovered in 1917 by Sturtevant as a suppressor of recombination between chromosomes in hybrids of different strains of Drosophila^1^. Inversions can be broadly classified on the basis by which they are formed as non-homologous end joining (NHEJ^2^), non allelic homologous recombination (NAHR) or fork stalling and template switching (FoSTeS^3^) inversions. NHEJ is a pathway for repairing double strand breaks in DNA. The inversion sequence ligates directly to break point without large homologous sequence^2^. NAHR is an aberrant recombination mechanism which occurs between homologous sequences. Homologous recombination between inverted repeats (IRs) will invert the intervening sequence and create an inversion^4^. Almost all (12/14) known large inversion (*>*1 Mb) polymorphisms are mediated by NAHR^5^. FoSTeS^3^ is a DNA replication error resulting in multiple copies of local sequences in both forward and reverse order. Although FoSTeS generates inverted sequences, we prefer to classify FoSTeS inversion as a type of complex copy number variation rather than a simple inversion.

Inversion polymorphisms remain one of the most poorly mapped classes of genetic variation. Before the advent of sequencing, it was only possible to identify large cytogenetically visible inversions via microscopy^6^. Inversions can be detected from aberrant linkage disequilibrium (LD) patterns from population single-nucleotide polymorphism (SNP) genotyping data, but this provides limited power to detect inversions smaller than 500 kb or with minor allele frequency less than 25%^7–9^. Inversions can be inferred from second generation sequence data by abnormal pair end mapping and split read alignment^10^. In theory this approach can be used to detect all NHEJ inversions^11^. Thus the remaining poorly understood inversions are NAHR inversions with a median size which cannot be detected using either a short read or cytogenetic approach. Third generation sequencing platforms, in particular Oxford Nanopore Technologies can sequence reads up to hundreds of kilobases, which is suitable to span IR in order to detect NAHR inversion. To fill the gap of poorly known inversion, we design a new tool, namely npInv (nanopore Inversion), for use with third generation sequencing data to detect long NAHR inversions.

## Results

### Detecting and genotyping inversion

We present npInv, a novel tool designed specifically for detecting and genotyping NAHR mediated inversions from long read sequencing data. The input to npInv is an alignment file in bam format generated from local aligner such as BWA-MEM^12^. npInv’s pipeline and pseudo code are shown in Figure 1 and in supplementary methods, respectively. In brief, npInv scans the alignment file for reads that contain pairs of subread alignments mapping to the same chromosome but with different orientation (Figure 2). npInv records this subread alignment pair as an inversion signal. If a subread alignment pair overlap in the original read, npInv records this overlapping sequence as an inverted repeat. npInv clusters and filters all the inversion signals in order to detect into inversion event based on position and the number of inversion signals in the cluster. npInv reports both the number of reads which support an inversion, as well as the number of reads supporting the non-inverted allele (reads which span the inversion breakpoints). Finally npInv applies a binomial model^13^ to genotype inversion from these read counts (see Methods). npInv reports the position, mechanism and genotype of each inversion.

**Figure 1.**
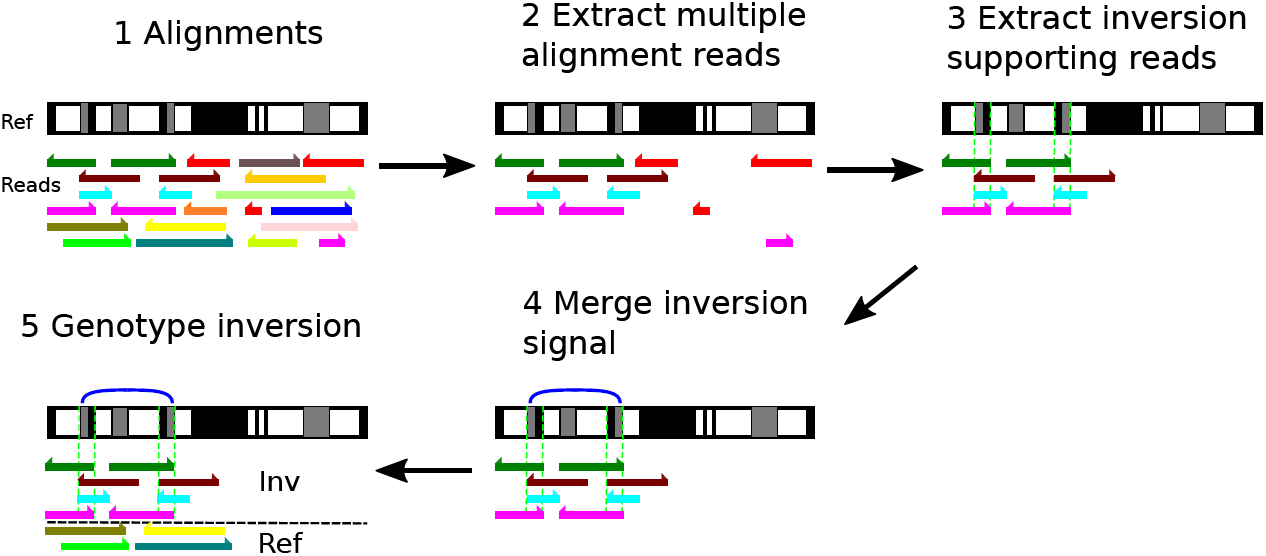
Software pipeline. The same colour bars indicate the alignment from the same reads. Half arrows indicate the orientation of the alignment. (1) The original alignment in a region. (2) Reads with multiple sub-alignments to the same chromosome are retained. Uniquely aligned reads are removed. (3) We obtain inversion signals and identify inverted repeats from sub-alignment. If inverted repeats (green dash lines) are observed, inversions are classified as NAHR, otherwise it is classified as NHEJ. Non inversion information reads are removed. (4) Inversion signals were merged into regions as blue arcs.(5) Once the inversion regions are defined, we estimated the number of inversion reads as well as the number of reads supporting the non-inverted (reference) allele, which are removed in the step (2). Horizontal black dash line indicates the classification of inversion and reference reads. Finally, the software predicts the inversion with position, mechanism and genotype.

**Figure 2.**
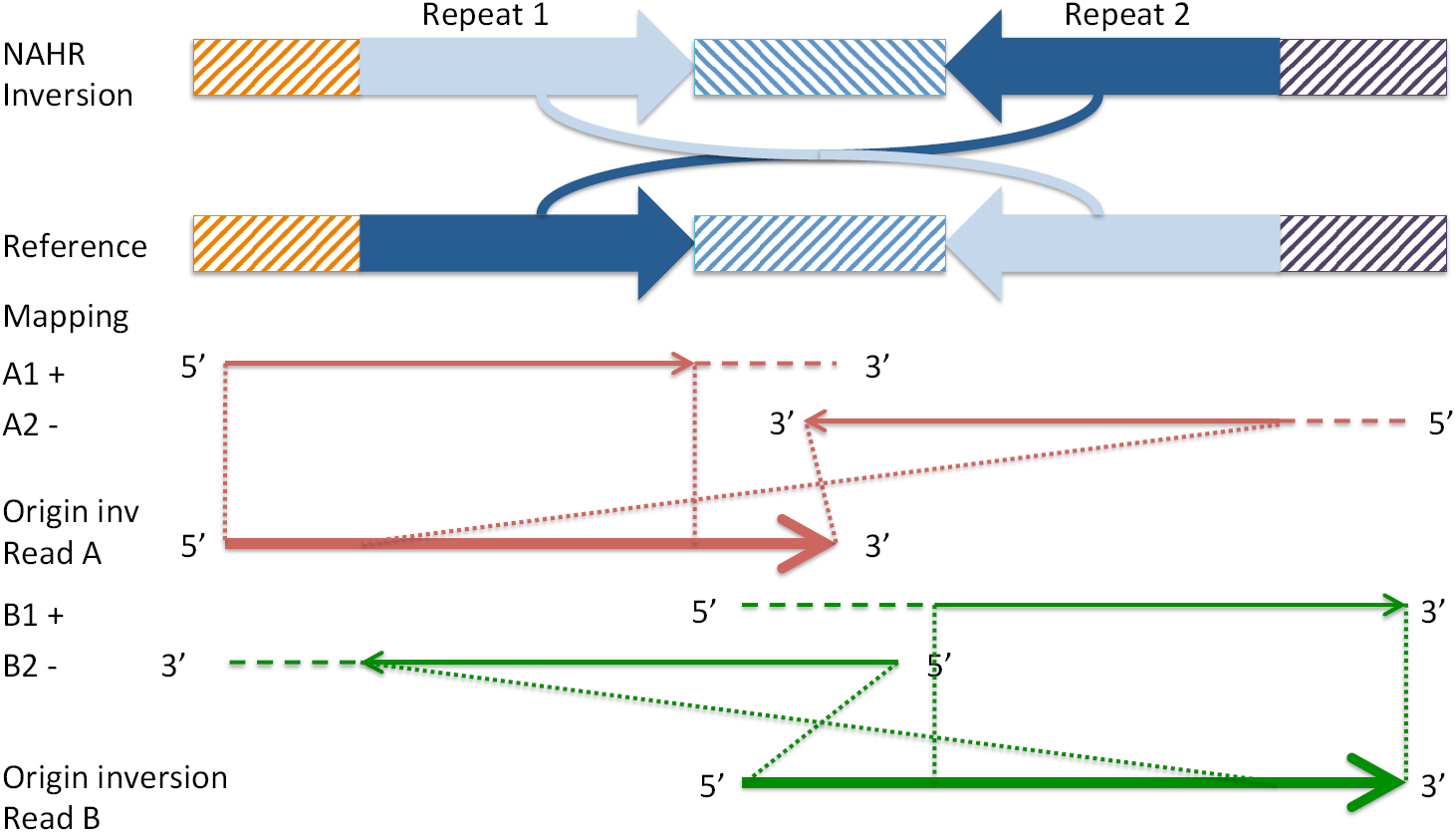
Illustration of effect of a NAHR mediated inversion on long read sub-alignments. Idealised NAHR inversion and reference are shown in first two panels. Inverted repeats are showed as dark and light blue. Orange, dark purple and blue hashed rectangles indicate unique sequence. The direction of the hashing indicates its orientation. The third panel (red) shows a read supporting the left breakpoint of the inversion. The large arrow indicates the original unmapped read. The smaller arrows indicate two sub-read alignments, with the direction of the arrow indicating the alignment orientation, and the horizontal dashed line indicating aligned and clipped sequence. The dot lines indicate the position of the subread alignment on the original read. The fourth panel (green) is similar to the third panel, except that it illustrates the read supporting the right breakpoint.

### Benchmarking the software using simulated data

We first benchmarked the software using simulation data. We simulated 61 NAHR, 100 short (*<*4 kb) and 100 long (*>*4 kb) NHEJ non-overlapping inversions in reference GRCh37 chromosome 21. NAHR inversions were simulated based on the location of IR of length above 500 bp in the reference chromosome 21 (which limited their number to 61). We randomly set the genotype of inversion to be heterozygous or homozygous. Next, we used readsim^14^ to simulate reads with an average read length of 3 kb, 6 kb or 9 kb. Sequence substitution, insertion and deletion rates were set at 5.1%, 4.9% and 7.8%, respectively based on previously described characteristics of nanopore sequence data^15^. Sequence depth was set at 5, 10, 20 or 40 fold for different simulations. Reads were aligned by BWA-MEM^12^. The alignment result was used for npInv, as well as for software

Lumpy^16^ and Sniffles^17^. The positive predictive value (PPV), sensitivity (S) and genotyping consistency (GC) were calculated for each datasets (Figure 3).

**Figure 3.**
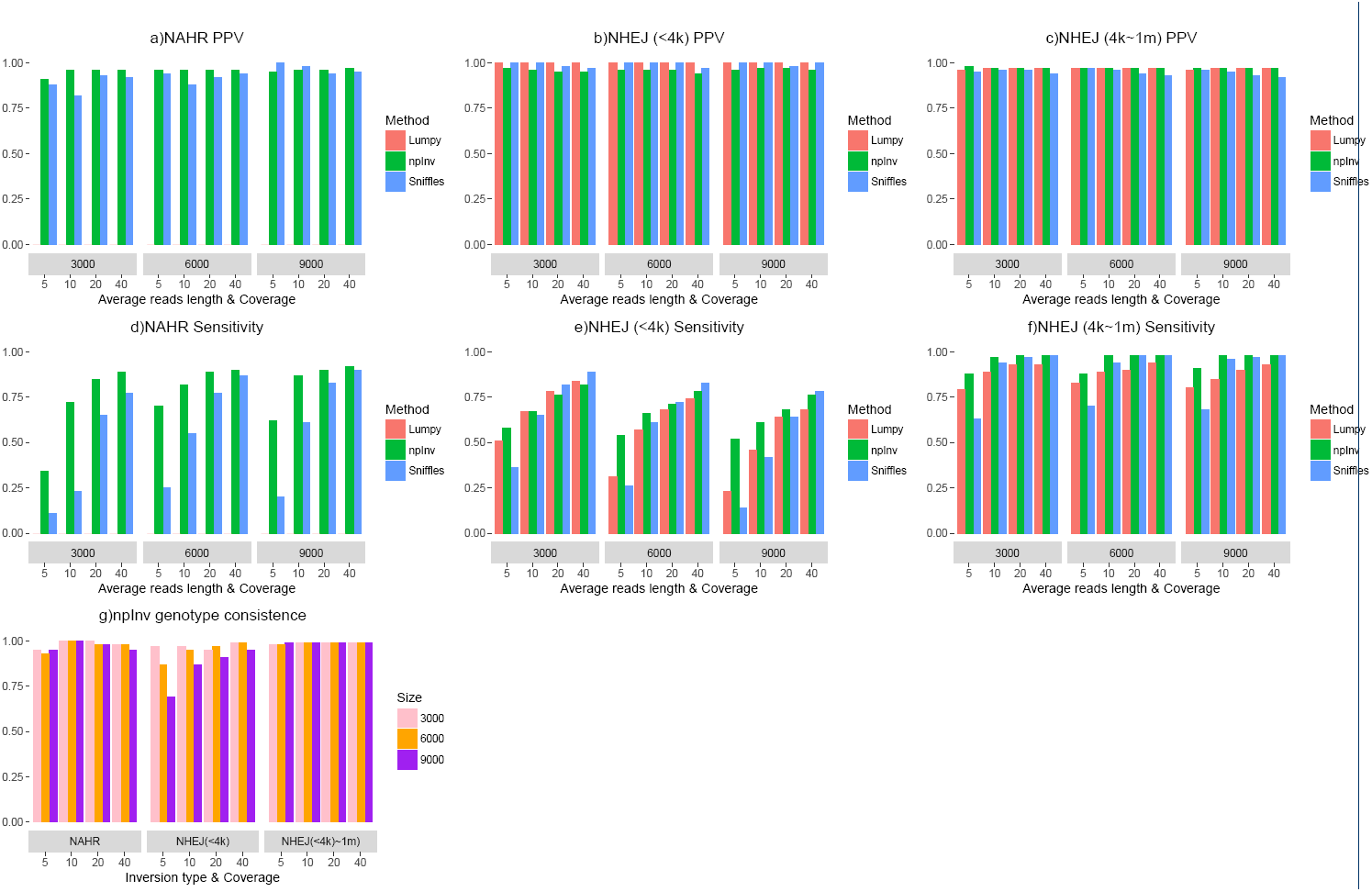
Performance comparison for npInv, Lumpy and Sniffles at three type of inversions. We simulated a diploid human chromosome 21 with three types of inversions: non-allelic homologous recombination (NAHR), non-homologous end joining (NHEJ) with size less than 4 kb and NHEJ with the size between 4 kb to 1 Mb (n = 61, 100 and 100, respectively). Software Lumpy^16^, Sniffles^17^ and npInv were applied to the above datasets. Positive predicted value (PPV) and sensitivity were estimated for each method. Lumpy did not detect NAHR inversion and genotype information is not available for Lumpy or Sniffles.

For simulated NAHR inversions, npInv demonstrated substantially better sensitivity (41% to 210%)than the next best program (Sniffles) over all coverage and read-lengths simulated (Fig, 3a). Lumpy didn’t predict any simulated NAHR inversions. Instead, it reported inversion breakpoints as potential structural variation. This was likely because its algorithm didn’t expect repetitive sequence around inversions. The PPV of npInv was also highest across most coverage and read-lengths, although Sniffles’ PPV, which was slightly (2% to 5%) higher than npInv in low coverage long read datasets (Fig, 3d). npInv’s PPV remained high (*>* 90%) across all datasets, while its sensitivity depended on the depth and read length. The sensitivity was good (*>* 80%) at 20 fold coverage and it did not improve significantly when the depth increased to 40 fold. The read length didn’t play a key role on both PPV and sensitivity, which was likely due to the fact that most of the background IR used to simulate NAHR inversions are of length less than the shortest average simulated read length (of 3 kb).

For NHEJ inversions the difference between the algorithms was not as pronounced. For long (*>* 4*kb*) NHEJ inversions, PPV for all 3 methods was more than 92%. The sensitivity of the three methods was similar (around 80%) for 20x coverage or higher, but npInv had a higher sensitivity at lower coverage. For short (*<*4 kb) NHEJ inversions, the PPV for all 3 methods was higher than 94%, but their sensitivity ranged from 26% to 89%. Lumpy’s sensitivity was lower than previously reported using simulations of highly accurate short paired-end reads^16^. For all 3 tools, the sensitivity decreased with increasing average read length. This reflects limitations of existing alignment algorithms on long error-prone reads. When the aligners align long read data, they have to decrease the penalty for gap opening and extending in order to adapt the relatively high sequencing error rate in long read sequencing. As a result, aligners preferred to incorrectly align more sequence at the inversion breakpoint. Even worse, when the inversion was short compared to the read length, the aligner might fully align the inversion spanning read to the reference with wrong gap opening and extending at the inversion flipping sequence (Figure 4). In this case, an inversion supporting read would be incorrectly regarded as a reference supporting read.

**Figure 4.**
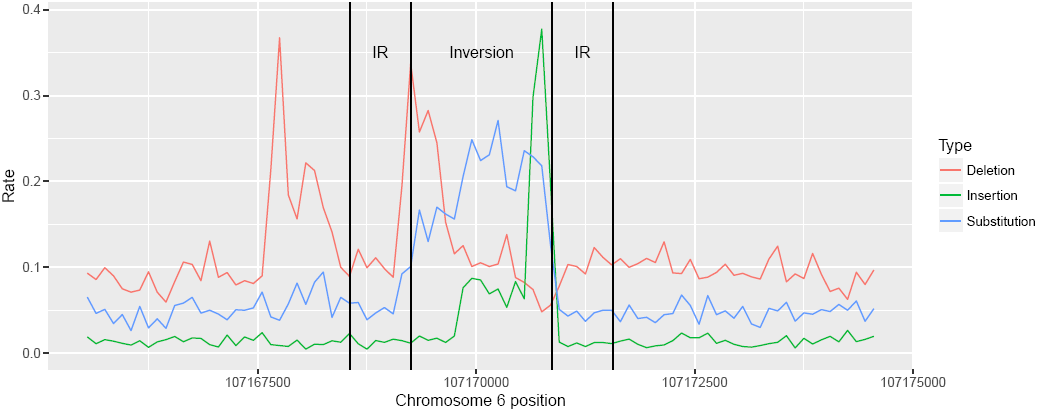
Error rates distribution around a wrong mapping homozygous inversion region. The border of inverted repeat (IR) and inversion are showed as vertical lines. Substitution, deletion and insertion rate are estimated from all the alignments in this region. Many alignments are spanning the whole inversion regions (figure not show). The high error rate within inversion indicates wrong alignments.

npInv was the only algorithm which reports the genotype for each inversion. To correctly genotype an inversion both the inversion read and reference read should be detected correctly. npInv’s genotype consistency was higher than 90% for long NHEJ inversions and NAHR inversions, but was lower for short NHEJ inversions with low coverage and long reads (9kb) (Fig, 3g). The genotyping error is mainly caused by the limits of sensitivity in detecting reads supporting the inversion, and as a result counting these reads as reference-supporting, leading to homozygous inversions being annotated as heterozygous. This was particularly a problem in conjunction with the issues regarding alignment to short inversions as discussed in the previous paragraph (Figure 5).

**Figure 5.**
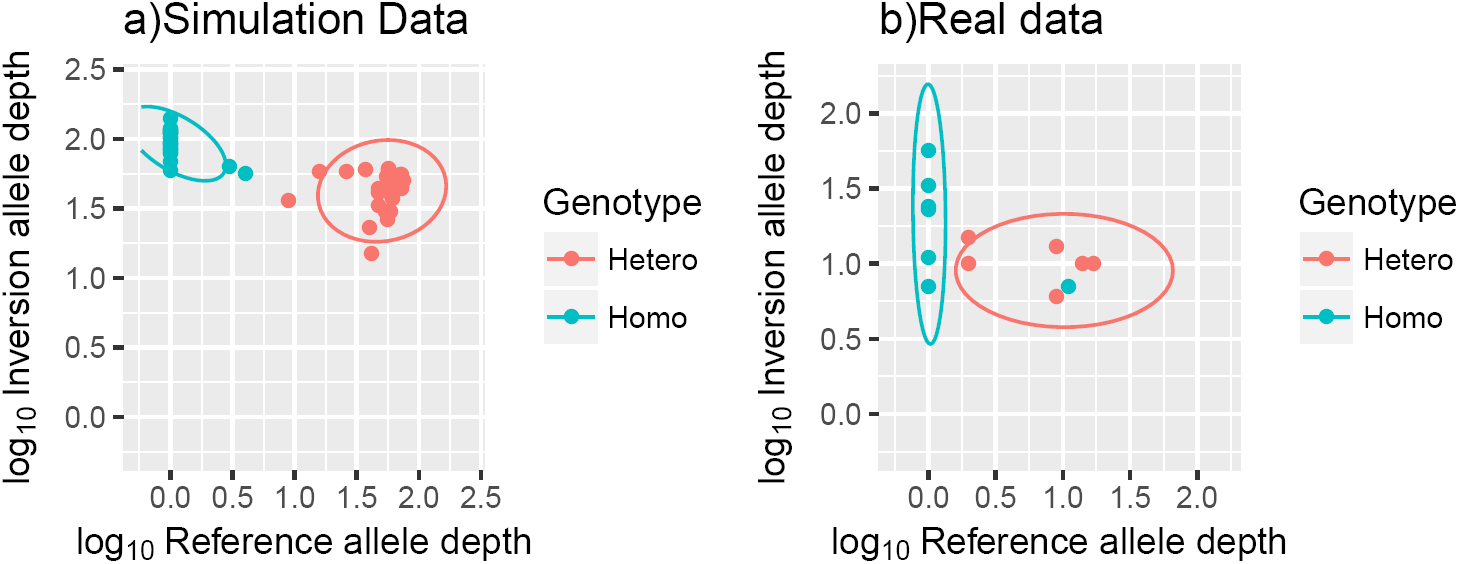
The performance of genotyping inversion from simulated and real data. Each dot is an inversion. X and Y axises are the base 10 logarithm of the estimated reference and inversion allele depth, respectively. “Hetero” and “Homo” are for heterozygous and homozygous inversion, respectively. Reference and inversion allele depth were estimated by npInv from simulation data (40 folds and average read length 6 kb NAHR inversion) and (b) real data (NA12878 nanopore data^18^). The genotype represented in the legend is the true genotype in the simulation (a) or validated database (b)^5^, respectively. The coloured ellipse indicates high confidence genotype prediction by npInv.

### Benchmarking the software using real data

We aligned Nanopore high coverage human sequencing data on sample NA12878^18^ to GRCh37 and identified 41 inversions using npInv. We compared our results to a ‘truth dataset’ of inversions from InvFest^5^, which is a database of validated inversions using various techniques including fluorescent in situ hybridization (FISH), polymerase chain reaction (PCR)^19–28^. We also compared this result to short read sequencing result of NA12878 by Delly^10^ and long read (Pacbio) assembly based on inversion call set^29^(Figure 6). npInv detected 18 (15 NAHR, 3 NHEJ) novel inversons and 23 (15 NAHR, 8 NHEJ) inversions overlapping one or more dataset, of which 13 inversions are included in the truth dataset. npInv analysis of nanopore sequence data had the largest overlap with the validated dataset compared to the PacBio assembly (5) and Delly Illumina analysis (8). This is because npInv (mean inversion size 61 kb) can detect both short and long inversions, while assembly (mean 1.8 kb) and Delly (mean 2.3 kb) preferentially identify short inversions. Inversions from the InvFEST database which could not be detected by npInv include inversions shorter than 2 kb (3), flanked by IR longer than 7 kb (5) or inversion with a deletion (1). In other words, npInv could detect all nine detectable (IR*<* 7kb) validated NAHR inversion as well as four out of five validated NHEJ inversion with size*>* 2kb.

**Figure 6.**
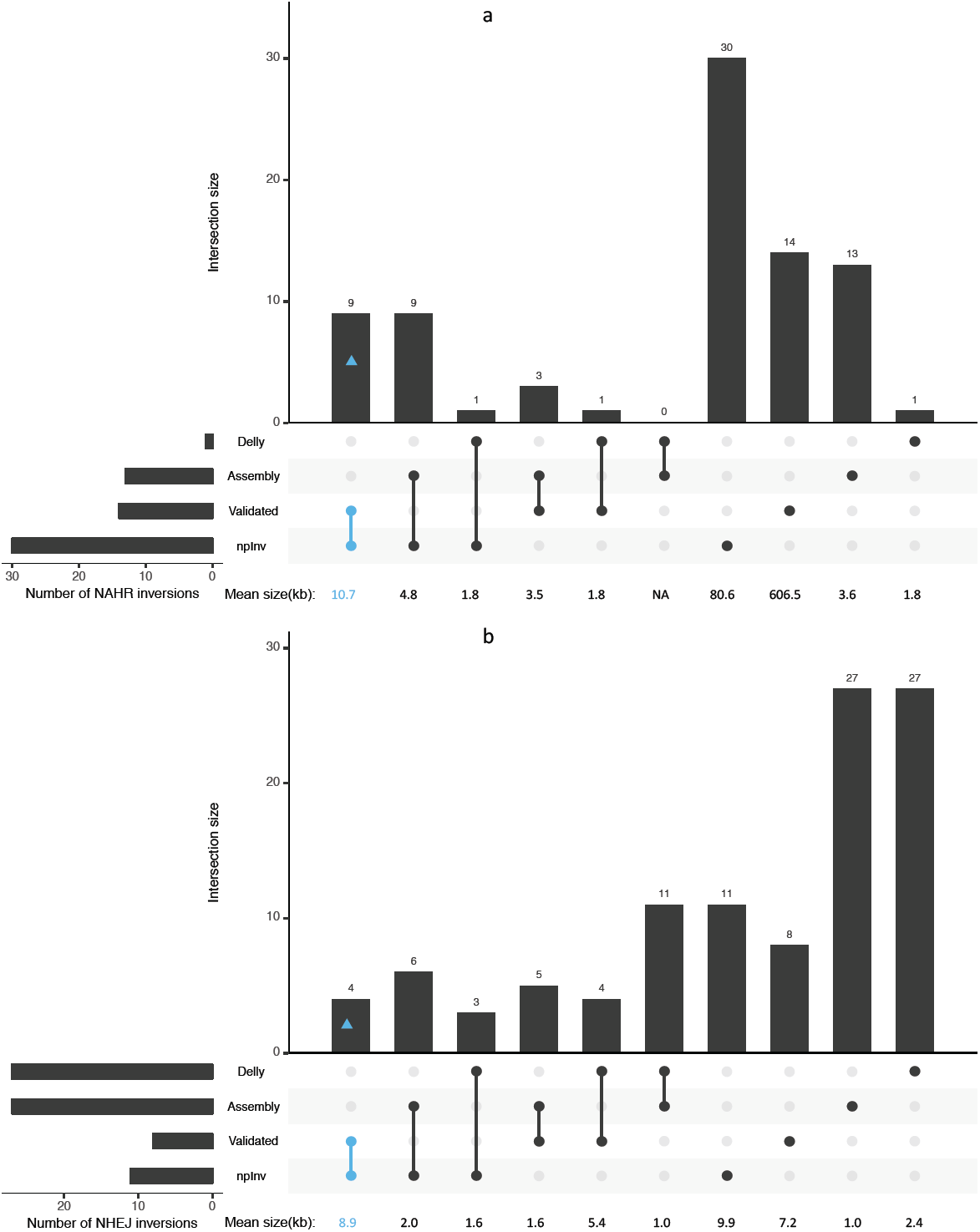
Intersection of four inversion datasets for individual NA12878. The four inversion datasets are labelled a. *Validated* (from InvFEST^5^); *Delly* (derived from Illumina sequence data by Sudmant, et al^30^); *Assembly* (derived from a PacBio assembly by Pendleton, et al^29^ and *npInv*, derived from nanopore sequence data. (a) and (b) are for NAHR and NHEJ inversions, respectively. The number of inversions in the intersection is shown in the bar chart. The connected dots below the bar chart indicate which methods are included in each intersection. The mean size of inversions in the intersection is shown under each bar. Intersection containing both npInv and validated are highlight with blue. The total number of predicted inversion is shown on the bottom left. This figure was generated using ggplot2**^?^** and modified version of UpSetR**^?^**.

We also used a set of validated 36 inversion sites in NA18278 (derived from InvFEST) to validate genotype consistency of npInv. The genotype consistency for homozygous reference, heterozygous inversion and homozygous inversion are 100%(23/23), 83%(5/6) and 86%(6/7) in the real data, respectively. Overall it is 94%(34/36).

### Experimental validation of novel inversions

We selected three novel inversions of size *>* 1kb predicted by npInv which could be validated using a PCR based approach. As this requires a PCR product which spans the inverted repeat, this placed an upper limit on the size of the IR to be less than 2 kb. Three of 18 novel inversions passed these criteria predicted from npInv. We checked inversion 4q35.2 (NHEJ), 3q21.3 (NAHR) and 10q11.22 (NHEJ) by PCR (Figure 7). Among these 3 inversions, there were 2 predicted heterozygous (4q35.2 and 3q21.3) and 1 homozygous inversion (10q11.22). We were able to validate predicted genotypes at both NAHR inversions. However, the 4q35.2 NHEJ inversion, could not be validated by PCR. Visual inspection of aligned nanopore reads revealed a clear structural variation breakpoint (Supplementary Figure 1a) which was also predicted to be an inversion by Sniffles^17^. However, inspection of Pacbio^29^ reads revealed almost no clipped reads, indicating an absence of an inversion (Supplementary Figure 1b). We surmise that the inversion observed in the nanopore sequence data may be due to a somatic mutation which occurred in a precursor cell to those used for Nanopore sequencing, however this is difficult to prove without access to the exact cell-line used in sequencing.

**Figure 7.**
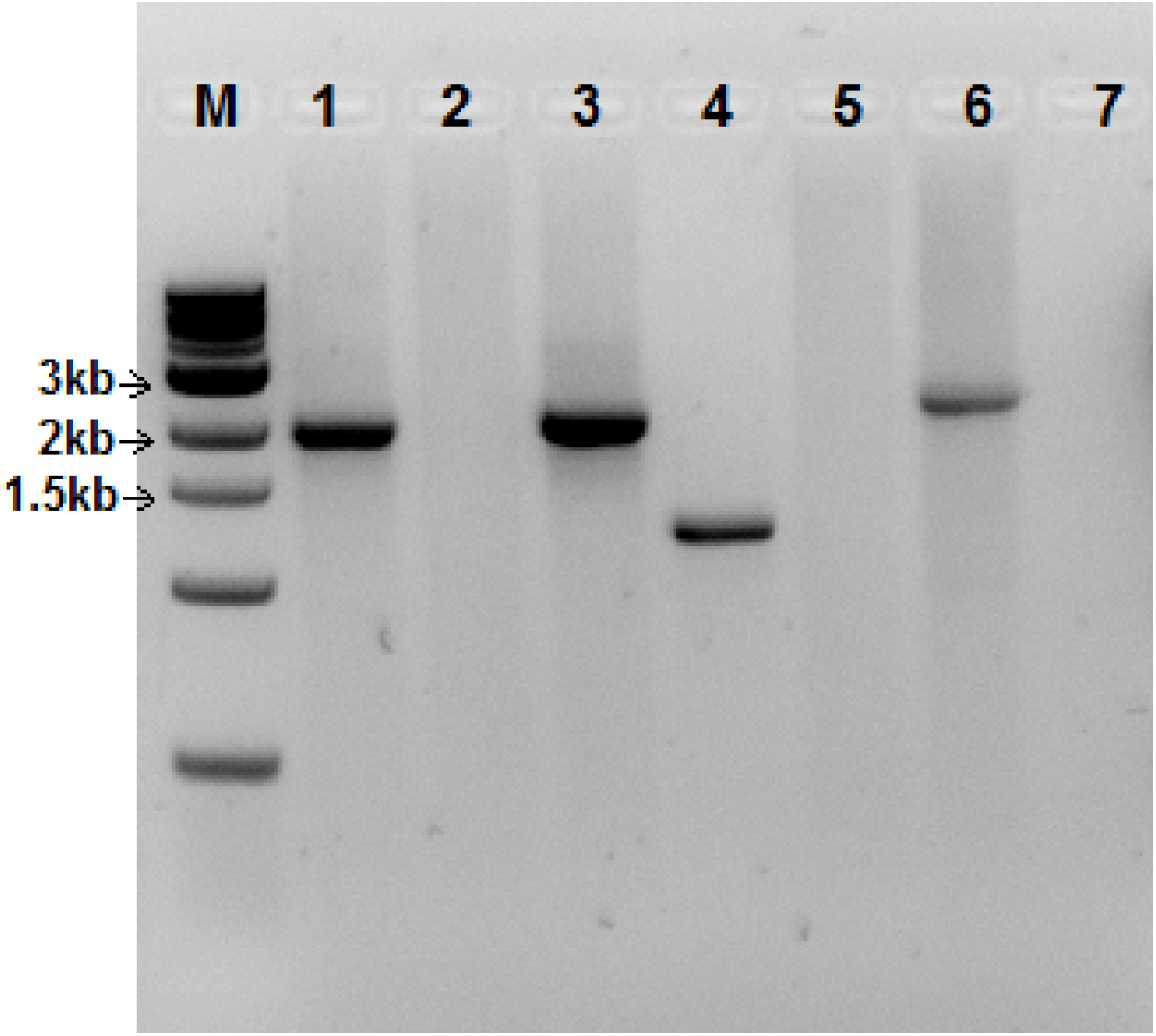
PCR products validating three inversions(4q35.2, 3q21.3 and 10q11.22). A band indicates this sequence exists in the genome. Lane1, 4q35.2 non-inverted. Lane2, 4q35.2 inverted. 4q35.2 is predicted to be heterozygous. Lane3, 3q21.3 non-inverted. Lane4, 3q21.3 inverted. 3q21.3 is predicted to be heterozygous. Lane5, 10q11.22 non-inverted. Lane6, 10q11.22 inverted. 10q11.22 is predicted to be homozygous. Lane7, Non template control. Primers could be found at supplementary material.

### Inversion map for NA12878

We combined all inversions detected by four different approaches on NA12878 including Delly applied to Illumina sequence data^30^, Pendleton et al. applied to Pacbio and Bionano sequence data^29^, InvFest database of validated inversions^5^, as well as novel inversions discovered by npInv. This resulted in a set of 87 known inversions, which we mapped to a karyogram ((Figure 8). We observed that NAHR inversions (mean size 275kbp) are longer than NHEJ or FoSTeS inversions (3.8 kb) (Figure 9)). Short read methods like Delly^10^ primarily focus on NHEJ inversion or NAHR inversion for which IR size is shorter than library insert size. Thus, it mainly reports the distribution of NHEJ inversions. On the other hand, the long read splitting method at IR like npInv could extend the range of detection to longer NAHR inversion(Figure 9).

**Figure 8.**
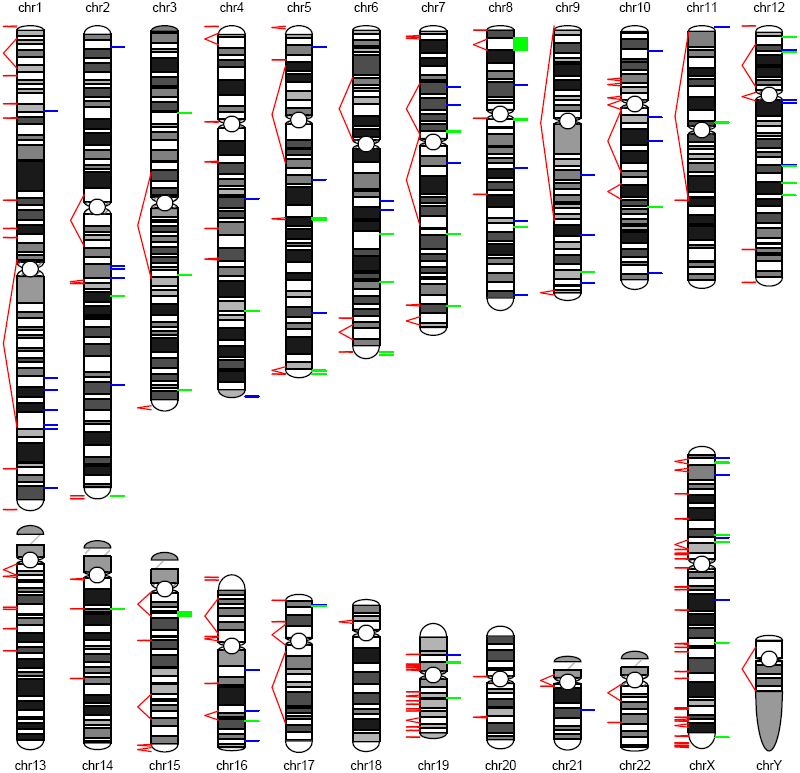
An inversion map for NA12878. A karyogram of human genome is depicted. The predicted (by existing methods) and potential (by genome compositions) inversions are shown on the right and left of the chromosome, respectively. Green bars are NAHR or palindrome inversions. Blue bars are NHEJ or FoSTeS inversions. Red line pairs indicate inverted repeat pairs which may mediate NAHR inversions. The bars and line pairs will be seen as a line if the distance between them is short.

**Figure 9.**
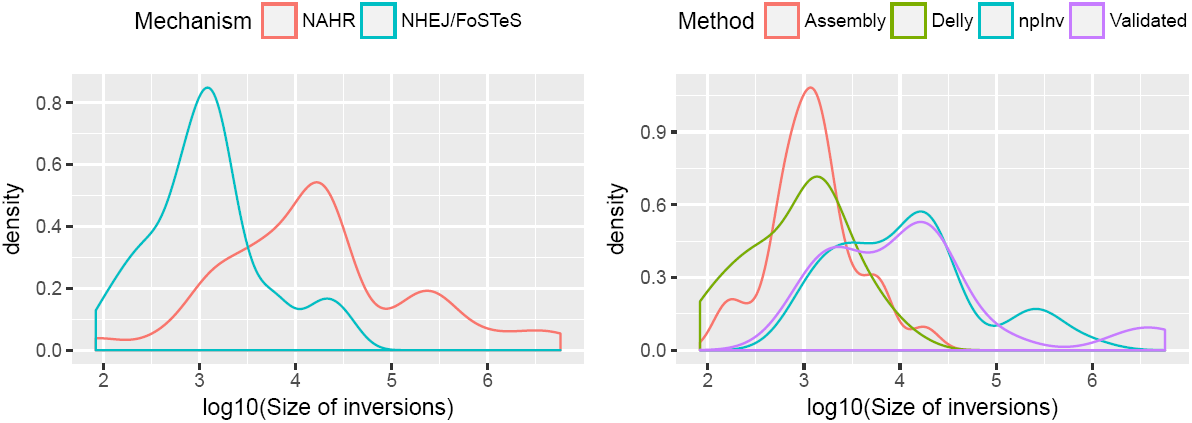
Inversion length distribution in density. a. Length distribution for NAHR and NHEJ/FoSTeS inversion. We broadly regard the non NAHR inversion as NHEJ/FoSTeS inversion. b. Length distribution for Delly^10, 30^, Assembly^29^, npInv and Validated^5, 19–28^ datasets. Density was estimated by function density in R.

We classified inversions according to the size of flanking IR as short (*<*500bp), median (500-7000bp) and long (*>*7kb). Short IR inversions can be detected by PEM based methods from short read sequencing data and local assembly^31^, particularly as the local sequence structure is typically not repetitive around short variants. Median IR inversions are efficiently detected using npInv as shown above.

### Characteristics of NAHR inversions

We investigated the relationship between IR and NAHR inversion by summarizing all the background IR in the genome as well as predicted and validated NAHR inversions (Figure 10, see Methods). The background IRs mainly occur with length less than 10 kb and between repeat distance ranging from 10 Mb to 100 Mb. There are two hotspots for IR at around 300bp and 6000bp, which is mainly due to the random distribution of short interspersed nuclear elements (SINEs) and long interspersed nuclear elements (LINEs) in the chromosome. If the probability of a NAHR inversion occurring is equal amongst all the IRs, the distribution of NAHR inversion should be the same as the distribution of background IR. However, we found the NAHR inversion distribution is totally different from the background IR distribution. Surprisingly, there is an almost linear relationship between the size of inverted repeat and the inversion, as well as an apparent empirical upper and lower bound on the size of an IR which can mediate an inversion of a certain size. For example, a 1Mb inversion can only be mediated by an IR of length greater than 50kb. This suggests only some IRs have the potential to mediate non-allelic homologous recombination and become a NAHR inversion. Furthermore, for small (size*<*3 kb) NAHR inversion, IR sequence must almost have a 100% identity. For larger (IR*>*50 kb) NAHR inversions, the size of first and second inverted repeat are not always the same and the identity could be lower (0.90 to 0.99). As the size of IR increase, the tolerance of recombination also increases.

**Figure 10.**
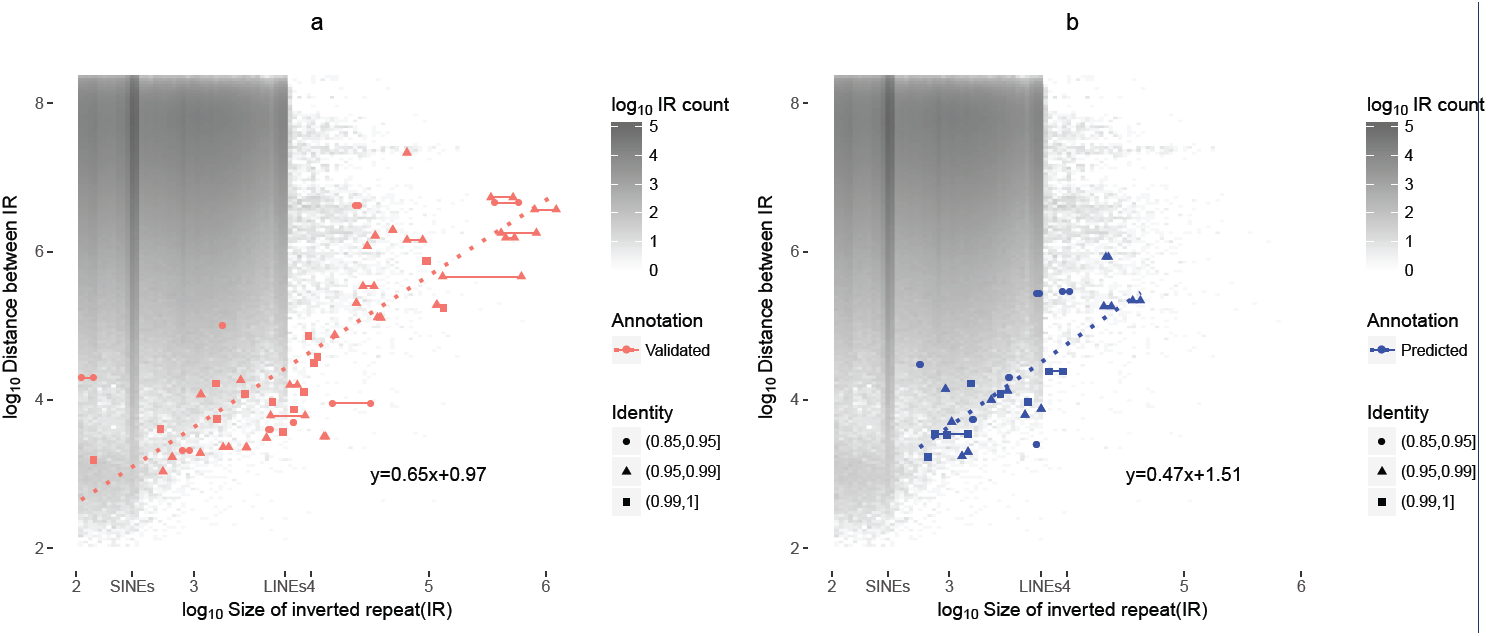
Relationship between inverted repeat (IR) length and distance between repeats. (a) is from all validated inversions from invFest and (b) is from predicted NAHR inversion at NA12878 using npInv. Two points are drawn for each pair of IR which mediates a NAHR inversion. In the majority of cases the pair of IRs have similar length and so the two points are co-localised; otherwise the two points are connected by a line. The y-axis indicates the distance between the IRs and the x axis indicates the length of the IR. The dotted line indicates the linear regression from log10 average size of IR to log10 size of inversion without flanking IR. The equation of it is shown at bottom right. The background heat map indicates the count of IR in the genome with given IR length and distance between IR. The size of short interspersed nuclear elements (SINEs, 300 bp) and long interspersed nuclear elements(LINEs, 7 kb) is shown in the x axis legend. Palindromic sequences are excluded.

We use this observed relationship to map the potential location of inversion mediated by large IR (*>*7kb) in the human genome, which are still not well characterized by sequencing based approaches. We filter all IR pairs with length greater than 7kb on the basis of the distance between the repeats (see Methods) to identify IR pairs which can mediate inversions (Figure 10). This leaves 140 regions in which large IR inversions could occur (Figure 8 and supplementary table). All of the 5 known NAHR inversions with IR greater than 7kb observed in NA18278 occur within one of these regions.

## Discussion and Conclusion

We developed a new tool, npInv, to detect and genotype inversion from long read sequencing data, with particular application to data generated using Oxford Nanopore sequencing technologies devices. The application of npInv shows high accuracy in both simulation and real data. We use npInv to uncover an almost linear relationship between inverted repeat and NAHR inversion and show the potential of providing an individual inversion map. With the possible widespread adoption of long read sequencing data, application of npInv could help extend our understanding of the extent of inversion polymorphism, their evolutionary significance and their clinical impact.

We report the most comprehensive whole-genome inversion map to date, consisting of 87 inversions, of which 38 are NAHR mediated inversions and the remained are NHEJ or FOSTES mediated. We exploited knowledge of potential sites of NAHR inversion to identify a further potential 140 inversion loci with IR length greater than 7kb. An increase in the yield of ultra-long (*>*100kb) sequence data on this sample, coupled with algorithmic improvements in alignment of long reads will help refine the location of inversions flanked by these long IR.

## Methods

### Inverted repeat mapping

Different size of invert repeats required different methods. Long (*>* 1*kb*) inverted repeat were identified by extracting inverted duplications from SD database^4^ at http://humanparalogy.gs.washington.edu/. This contains long invert repeats with long (*>* 10*kb*) insertions or deletions. Median(*>* 500*bp*) inverted repeat were identified by running inverted repeat finder^32^(IRF, version 3.05) for each chromosome. The parameter was 2 3 5 80 10 800 50000 300000 -d -h -t4 1000 -t5 10000 -t7 300000. Short (*>* 100*bp*) inverted repeats were identified by using last^33^(v458) to align each chromosome to its self. Each alignment pair with identity greater than 0.90 was defined as an inverted repeat pair. The parameter was -s 0 (reversed alignment).

For figure 8, we first filtered the IR (inferred from SD database^4^) less than 7k to identify 2130 IRs. We carried out a linear regression of IR length against the inversion size on all InvFEST and all npInv detected inversions in NA12878 respectively. The regression parameters obtained are similar. We use the InvFEST regression parameters to build a predictive model of the length of IR against the distance between IR (i.e. the minimum potential inversion size). We removed the IRs outside of 90% prediction interval by R to identify 1302 IRs. Then we applied BEDTools^34^ to sort and merge these regions into 140 non-overlapping genomic regions.

### Analysis of long read sub-alignment

npInv focuses on reads which have multiple sub-alignments. For each of these sub-read alignments, i, we sort the alignments by its left-most location in the read. Then we record the 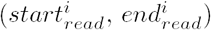 co-ordinates of the alignment on the read, the 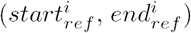 co-ordinates in the reference genome, as well as the reference orientation and chromosome. For a read containing multiple sub-alignments (Figure 2), we perform the following analysis. We first filter alignments with length less than 500 bps or for which the alignment interval on the read is totally contained by another alignment interval. Next, for each pair of read-adjacent alignment intervals (which are allowed to overlap), we keep pairs mapping to the same chromosome and in different alignment orientation as potential inversion signals (A1-A2, B1-B2 in Figure 2). If the first sub alignment is in forward strand, we record this signal as an inversion forward supporting signal. Otherwise, we record as reverse signal. If the first alignment’s location in reference is bigger than the second alignment, we record this signal as a left breakpoint inversion supporting signal. Otherwise, we record this signal as right. If two alignment intervals are overlapping by more than 500bp (on the read), this inversion signal is considered to be mediated by NAHR. The overlapping sequence could align to the inverted repeat in both orientations (light blue and dark blue in Figure 2). Only one of these pair of read alignments includes sequence from the inversion itself (Figure 2). All inversion signals are sorted by chromosome and left-most start position on reference.

### Analysis of inversion signal

After scanning the bam file by split read analysis, the software identifies numerous inversion signals. The user has the option of providing a database of known IR pairs in the genome. If this is provided, the software creates a bin for each IR in the database which is used to merge inversion signals. Each bin represents an inversion call. For each candidate inversion, we check whether the left breakpoint and right breakpoint are within X bp (in practice, X=2000) in the IR database’s left and right repeat sequence. If true, group them in this IR bin and delete the binned signal. If the user does not provide an IR database, and also for the remaining signals which cannot be clustered around known IR, inversion signals are grouped into the same bin if their reference start and end are both less than X bp (default 2000) from each other. We then investigate whether each merged inversion signal contains supporting reads on both forward and reverse strands, and also at both left and right breakpoints. The output inversion’s start and end are the mean value of the left breakpoints and right breakpoints, respectively. The output inversion left and right breakpoint start and end are the minimum and maximum alignment position at left and right breakpoint in the reference, respectively. We calculate the inversion supporting read *R*_*inv*_ as the sum of reads supporting the left and right breakpoint signal for genotyping inversion. If a read supports both left and right breakpoint, it will count as one left breakpoint signal and one right breakpoint signal.

npInv annotates the inversion longer than *L*(parameter, default 1M) as long inversion. We consider that long inversion is not reliable for either NHEJ (usually shorter than 1 Mb) or NAHR (likely with inverted repeat which is too long to be fully spanned) inversion.

### Analysis of non inversion signal

We calculate the average substitution, deletion and insertion rate and their standard deviations from the first min(10000*, all*) primary alignments. For each alignment overlapping with inversion, we define its inversion region as (max(left breakpoint start, alignment start), min(right breakpoint end, alignment end)). We calculate three error rates (substitution, deletion and insertion rate) in its inversion region from the primary alignments. If the all three error rates are less than its average rate plus its one standard deviation, we kept this alignment as a reference supporting alignment. We calculate the sum of reference supporting read *R*_*re f*_ for the next step. If a read spans both left and right breakpoint and passes the criteria, it will count as one reference left breakpoint signal and one reference right breakpoint signal.

### Inversion genotyping

For each binned inversion, we get the number of inversion and reference supporting reads (*R*_*inv*_,*R*_*re f*_) from the above step. Applying binomial model^13^ on the genotyping inversion, the posterior probability *P* of genotype *G* = *g*_*re f re f*_ *, g*_*re f inv*_*, g*_*invInv*_ conditional on the observed read counts *R*_*re f*_ and *R*_*inv*_ could be written as below.

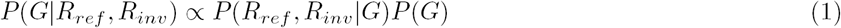

The likelihood *P*(*R*_*re f*_ *, R*_*inv*_*|G*) could be written as

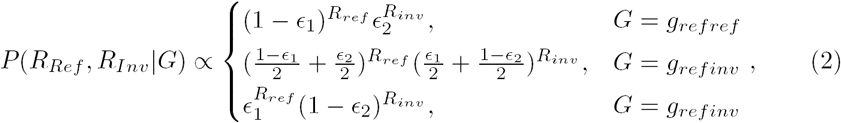

where *ε*_1_ and *ε*_2_ are the error rates of incorrectly assigning an inversion-supporting read to a reference supporting read and vice-versa, respectively(in practice, we use *ε*_1_ = *ε*_2_ = 0.01, however with availability of more data it would be possible to infer specific mis-assignment rates). We assume an uniform prior such that *P*(*G* = *g*_*inv*_) = *P*(*G* = *g*_*re f*_) = 0.5 and then by Hardy-Weinberg equilibrium *P*(*G* = *g*_*re f re f*_) = *P*(*G* = *g*_*invInv*_) = 0.25 and *P*(*G* = *g*_*re f inv*_) = 0.5. Then we choose the maximum posterior probability genotype as the genotype for the individual. The genotype quality Q is calculated as the second maximum posterior probability *P*_2*nd*_ divided by the maximum posterior probability *P*_1*st*_ in Phred quality score as below.

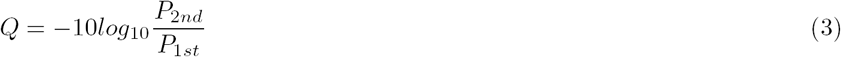

### Inversion simulation and benchmarking

We chose the whole GRCh37 chromosome 21 as the reference. We grouped the inversions into three types, which were NAHR, short (0-4 kb) NHEJ and long (4 kb to 1 Mb) NHEJ inversions. We simulated 61 NAHR, 100 short and 100 long NHEJ non-overlapping inversions in reference chromosome 21. NAHR inversions were simulated based on the reference IR (*>*500 bp) from IRF^32^ and limited to 61 non-overlapping NAHR inversions on chromosome 21. We randomly set the genotype of inversion as heterozygous or homozygous. Then we simulated a diploid chromosome 21 and flipped over the simulated inversion interval in one or two chromosomes according to its genotype. Next, we used readsim^14^(version 1.6) to simulate reads from this diploid chromosome with an average read length of 3 kb, 6 kb or 9 kb. Sequence substitution, insertion and deletion rates were set at 5.1%, 4.9% and 7.8%, respectively based on previously described characteristics of nanopore sequence data^15^. Sequence depth was set at 5, 10, 20 or 40 folds for different simulations. The readsim parameter is sim fa –rev strd on –tech nanopore –read mu 3000,6000,9000 –read dist exp –cov mu 5,10,20,40 –err sub mu 0.051 –err in mu 0.049 –err del mu 0.078.

Simulation reads were aligned by BWA-MEM^12^(version 0.7.15-r1142-dirty) to chromosome 21. The BWA-MEM parameter is -t 16 -x ont2d -M, which is suggested by Sniffles’ readme. The alignment result was used for npInv (version 1.2), as well as for software Lumpy^16^ and Sniffles^17^. We run Lumpy (v0.2.13) from its executable file named lumpy with parameter -mw 4 -tt 1e-3 -sr bam file:BAMINPUT,back distance:20,weight:1,id:1,min mapping threshold:1. Sniffles (version 1.0.5) was downloaded from https://github.com/fritzsedlazeck/Sniffles. We applied Sniffles directly to the simulation bam files with the default parameter. Lumpy or Sniffles inversions were called when their vcf ALT fields are equal to *<*INV*>*. An inversion was classified as positive predictive inversion when the true simulation inversion interval was 90% overlapping with the predictive inversion interval, and vice-versa. Finally, the positive predictive value (PPV), sensitivity (S) and genotype consistency (GC) were calculated for different datasets.

### Detection of inversion on NA12878

NA12878 raw data^18^(version rel3) was downloaded from https://github.com/nanopore-wgs-consortium/ NA12878. We aligned it to GRCh37 by BWA-MEM^12^(version 0.7.15-r1142-dirty). The key parameter was -k11 -W20 -r10 -A1 -B1 -O1 -E1 -L0 -Y. We ran npInv(version 1.2) with default parameter. The predicted inversions are the inversions whose “FILTER” field is equal to “PASS” in vcf^35^ file.

### Classification of inversion by mechanism

For inversions from InvFEST, we accepted the mechanism from InvFEST. For the remaining unclassified inversions, we checked whether the start and end were within inverted repeats from the SD database^4^. If an inverted repeat was found, we classified the inversion as NAHR mediated with IR sizes and identity from SD database. Otherwise, we extracted the whole inversion sequence and aligned it to itself by YASS^36^. If the YASS’s dotplot showed inverted repeat sequence at both the start and the end, we classified it into NAHR inversion. The IR sizes and identity were determined by the YASS’s alignment result. When the inversion was totally reverse complement, we classified it as Palindrome. We classified the remaining inversions into NHEJ/FoSTeS inversion.

### PCR validation

PCR was used to validate 3 inversions detected from the sequencing data. Two forward primers were designed to overlap the inversion breakpoints, one to amplify the reference copy and a second primer to amplify the inverted copy with a shared reverse primer. PCR reactions were performed using 1x HotStar Taq DNA Polymerase (Qiagen), 2.5mM MgCl_2_, 200nM of forward primer (either to amplify the reference or the inverted sequence), 200nM reverse primer and 2ng/uL of DNA NA12878. PCR conditions were optimized for each PCR target. The following PCR conditions were used: hot start at 95*°*C for 15 minutes, 35 cycles of 95*°*C for 30s, 60*°*C for 30s and 72*°*C for 4 minutes with a final extension of 10 minutes at 72*°*C. An annealing temperature of 55*°*C was used to amplify the inverted sequence of 3q21.3. PCR products were analyzed by horizontal electrophoresis on 1.5% agarose gel.

## Declarations

### Ethics approval and consent to participate

Not applicable.

### Consent for publication

Not applicable.

### Availability of data and materials

npInv(version 1.2) is an open-source java runnable jar file available at https://github.com/haojingshao/npInv with license GPL-3.0. It was run and tested on Mac OS and Linux with java version 1.8.0 66 and 1.8.0 112.

### Authors’ contributions

L.C. and C.H conceived the study. H.S. and M.C. performed the analysis. H.S developed the software. D.G and T.D. performed the experiment. H.S. wrote the manuscript, which was revised and approved by all authors.

## Acknowledgements

We would like to thank Jain et al.^18^ for nanopore open dataset.

## References

1. Sturtevant, A. H. Genetic factors affecting the strength of linkage in drosophila. Proceedings of the National Academy of Sciences 3, 555–558 (1917).

2. McVey, M. & Lee, S. E. Mmej repair of double-strand breaks (director’s cut): deleted sequences and alternative endings. Trends in Genetics 24, 529–538 (2008).

3. Zhang, F. et al. The dna replication fostes/mmbir mechanism can generate genomic, genic and exonic complex rearrangements in humans. Nature genetics 41, 849–853 (2009).

4. Bailey, J. A. & Eichler, E. E. Primate segmental duplications: crucibles of evolution, diversity and disease. Nature Reviews Genetics 7, 552–564 (2006).

5. Martínez-Fundichely, A. et al. Invfest, a database integrating information of polymorphic inversions in the human genome. Nucleic acids research gkt1122 (2013).

6. Feuk, L., Carson, A. R. & Scherer, S. W. Structural variation in the human genome. Nature Reviews Genetics 7, 85–97 (2006).

7. Bansal, V., Bashir, A. & Bafna, V. Evidence for large inversion polymorphisms in the human genome from hapmap data. Genome research 17, 219–230 (2007).

8. Cáceres, A., Sindi, S. S., Raphael, B. J., Cáceres, M. & González, J. R. Identification of polymorphic inversions from genotypes. BMC bioinformatics 13, 28 (2012).

9. Sindi, S. S. & Raphael, B. J. Identification and frequency estimation of inversion polymorphisms from haplotype data. Journal of computational biology 17, 517–531 (2010).

10. Rausch, T. et al. Delly: structural variant discovery by integrated paired-end and split-read analysis. Bioinformatics 28, i333–i339 (2012).

11. Lledó, J. I. L. & Cáceres, M. On the power and the systematic biases of the detection of chromosomal inversions by paired-end genome sequencing. PLoS One 8, e61292 (2013).

12. Li, H. Aligning sequence reads, clone sequences and assembly contigs with bwa-mem. arXiv preprint arXiv:1303.3997 (2013).

13. Shao, H. et al. A population model for genotyping indels from next-generation sequence data. Nucleic acids research gks1143 (2012).

14. Schmid, R., Schuster, S., Steel, M. & Huson, D. Readsim-a simulator for sanger and 454 sequencing. View Article PubMed/NCBI Google Scholar (2006).

15. Jain, M. et al. Improved data analysis for the minion nanopore sequencer. Nature methods 12, 351–356 (2015).

16. Layer, R. M., Chiang, C., Quinlan, A. R. & Hall, I. M. Lumpy: a probabilistic framework for structural variant discovery. Genome biology 15, R84 (2014).

17. Sedlazeck, F. Sniffles. https://github.com/fritzsedlazeck/Sniffles (2016).

18. Jain, M. et al. Nanopore sequencing and assembly of a human genome with ultra-long reads. bioRxiv 128835 (2017).

19. Korbel, J. O. et al. Paired-end mapping reveals extensive structural variation in the human genome. Science 318, 420–426 (2007).

20. Kidd, J. M. et al. Mapping and sequencing of structural variation from eight human genomes. Nature 453, 56–64 (2008).

21. Pang, A. W. et al. Towards a comprehensive structural variation map of an individual human genome. Genome biology 11, R52 (2010).

22. Wang, J. et al. The diploid genome sequence of an asian individual. Nature 456, 60–65 (2008).

23. Ahn, S.-M. et al. The first korean genome sequence and analysis: full genome sequencing for a socio-ethnic group. Genome research 19, 1622–1629 (2009).

24. McKernan, K. J. et al. Sequence and structural variation in a human genome uncovered by short-read, massively parallel ligation sequencing using two-base encoding. Genome research 19, 1527–1541 (2009).

25. Stefansson, H. et al. A common inversion under selection in europeans. Nature genetics 37, 129–137 (2005).

26. Giglio, S. et al. Heterozygous submicroscopic inversions involving olfactory receptor–gene clusters mediate the recurrent t (4; 8)(p16; p23) translocation. The American Journal of Human Genetics 71, 276–285 (2002).

27. Osborne, L. R. et al. A 1.5 million–base pair inversion polymorphism in families with williams-beuren syndrome. Nature genetics 29, 321–325 (2001).

28. Gimelli, G. et al. Genomic inversions of human chromosome 15q11–q13 in mothers of angelman syndrome patients with class ii (bp2/3) deletions. Human molecular genetics 12, 849–858 (2003).

29. Pendleton, M. et al. Assembly and diploid architecture of an individual human genome via single-molecule technologies. Nature methods 12, 780–786 (2015).

30. Sudmant, P. H. et al. An integrated map of structural variation in 2,504 human genomes. Nature 526, 75–81 (2015).

31. Treangen, T. J. & Salzberg, S. L. Repetitive dna and next-generation sequencing: computational challenges and solutions. Nature Reviews Genetics 13, 36–46 (2012).

32. Warburton, P. E., Giordano, J., Cheung, F., Gelfand, Y. & Benson, G. Inverted repeat structure of the human genome: the x-chromosome contains a preponderance of large, highly homologous inverted repeats that contain testes genes. Genome research 14, 1861–1869 (2004).

33. Kielbasa, S. M., Wan, R., Sato, K., Horton, P. & Frith, M. C. Adaptive seeds tame genomic sequence comparison. Genome research 21, 487–493 (2011).

34. Quinlan, A. R. Bedtools: the swiss-army tool for genome feature analysis. Current protocols in bioinformatics 11–12 (2014).

35. Danecek, P. et al. The variant call format and vcftools. Bioinformatics 27, 2156–2158 (2011).

36. Noé, L. & Kucherov, G. Yass: enhancing the sensitivity of dna similarity search. Nucleic acids research 33, W540–W543 (2005).

